# Sowing date effects on anther dehiscence, pollen germination on the stigma, and fertility under heat in Japanese rice

**DOI:** 10.64898/2026.03.17.712342

**Authors:** Kazuto Kimura, Tomoaki Yamaguchi, Tsutomu Matsui

**Author notes:** Email addresses: Kazuto Kimura Tomoaki Yamaguchi Tsutomu Matsui.

## Abstract

Heat-tolerant rice cultivars are essential for mitigating global warming impacts. Basal anther dehiscence length (BDL) is a promising visible morphological marker for heat tolerance through stable pollination. We investigated the effects of sowing date on anther morphology, pollination, and fertility under controlled high-temperature conditions (35, 37, or 39 °C at flowering). Three *japonica* cultivars—‘Akitakomachi’ (early heading), ‘Koshihikari’ (medium), and ‘Hatsushimo’ (late)—were sown monthly over 3 months and grown in pots. At heading, the plants were exposed to the temperature treatments for 3 days, and the proportion of florets with ≥10 germinated pollen grains on the stigma (GP10) and seed set were assessed. Among anther traits, BDL showed the greatest variation, with all cultivars from the second sowing exhibiting the shortest BDL. Analysis of variance revealed significant effects of genotype, sowing date, and their interaction on anther traits and fertility. Regression analysis indicated that fertility was associated with GP10, with BDL contributing significantly to GP10 in the late-heading ‘Hatsushimo’, together with maximum temperature at flowering. Thus, both genotype and environment shape anther morphology, pollination, and fertility, indicating that BDL plasticity and genotype-specific environmental responses must be carefully considered when using BDL as a breeding marker for heat tolerance.

**Highlight:** Variation in sowing date significantly affects anther morphology and heat tolerance in rice. Genotype-specific responses to the growing environment require careful consideration for reliable breeding assessments.

## Introduction

Global warming is projected to exert profound impacts on agricultural productivity worldwide (IPCC, 2022). Rising temperatures have become a critical concern for staple crops owing to detrimental effects on yield (Zhao *et al*., 2017). In rice, simulation models based on controlled-environment data predict that global warming will increase the frequency of heat-induced floret sterility across Asia (Horie, 2019). Field observations of high-temperature–induced floret sterility support these projections (Osada *et al*., 1973; Matsushima *et al*., 1982; Matsui *et al*., 2021; Ishimaru *et al*., 2022; Matsui *et al*., 2025).

The tolerance of rice cultivars to heat-induced sterility can vary by more than 3 °C (Satake and Yoshida, 1978). Consequently, the development and adoption of heat-tolerant cultivars represents an effective strategy for mitigating yield losses under global warming (Nakagawa *et al*., 2004). Stable pollination is an important trait underlying such tolerance under field conditions (Matsui *et al*., 2021). During anthesis, pollen grains are released through dehiscences that form at both the apical and basal parts of the theca. A long basal dehiscence promotes stable pollination (Matsui and Kagata, 2003), even at high temperatures (Matsui *et al*., 2005). Because dehiscence size is genetically controlled (Tazib *et al*., 2015; Zhao *et al*., 2016), long basal dehiscence may serve as a simple, visible, and practical morphological marker for screening rice genotypes with high-temperature tolerance.

However, environmental factors such as photoperiod may influence anther dehiscence and heat tolerance. Early studies in cultivars of *Oryza sativa* L. suggested that short-day treatments disrupted dehiscence (Yasuda, 1944). More recently, Okamoto and Hara (2021) showed that short-day treatments applied around the panicle initiation stage shortened anther length, reduced pollen production, and increased floret sterility in *O. sativa* ‘Shinrei’. In contrast, Tanno *et al*. (2002) reported that short-day treatments that shortened the time to heading increased seed set across 17 *O. sativa* L. cultivars on average. If day length affects anther morphology and dehiscence ability, variations in sowing date or latitude between tropical and temperate environments may also exert similar effects, potentially leading to misjudgment of heat tolerance during cultivar screening.

Here we elucidated how shifts in sowing date (which encompasses changes in environmental conditions, including day length) affect anther dehiscence and tolerance to heat-induced sterility in *O. sativa* cultivars outdoor. Three cultivars with contrasting heading earliness were sown on three staggered dates and exposed to three high-temperature regimes during flowering. Anther dehiscence, pollination behavior, and seed set were analyzed to clarify the mechanisms underlying cultivar differences in heat tolerance.

## Materials and Methods

### Plant materials

Three *O. sativa* cultivars with different heading earliness—‘Akitakomachi’ (early), ‘Koshihikari’ (medium), and ‘Hatsushimo’ (late)—were selected from the prefectural government-recommended cultivars (Gifu Prefecture, Japan). Seeds were sown on 15 April, 15 May, and 25 June 2024 (Table 1). After puddling, seedlings were transplanted at the 5½-leaf stage into 1/10 000-a pots in a circular pattern, at 10 plants per pot. Each pot contained 1.2 kg of air-dried sandy loam soil (pH 7.64). A slow-release compound fertilizer was applied before puddling at rates equivalent to 0.26 g N, 0.17 g P□O□, and 0.17 g K□O per pot.

**Table 1.**
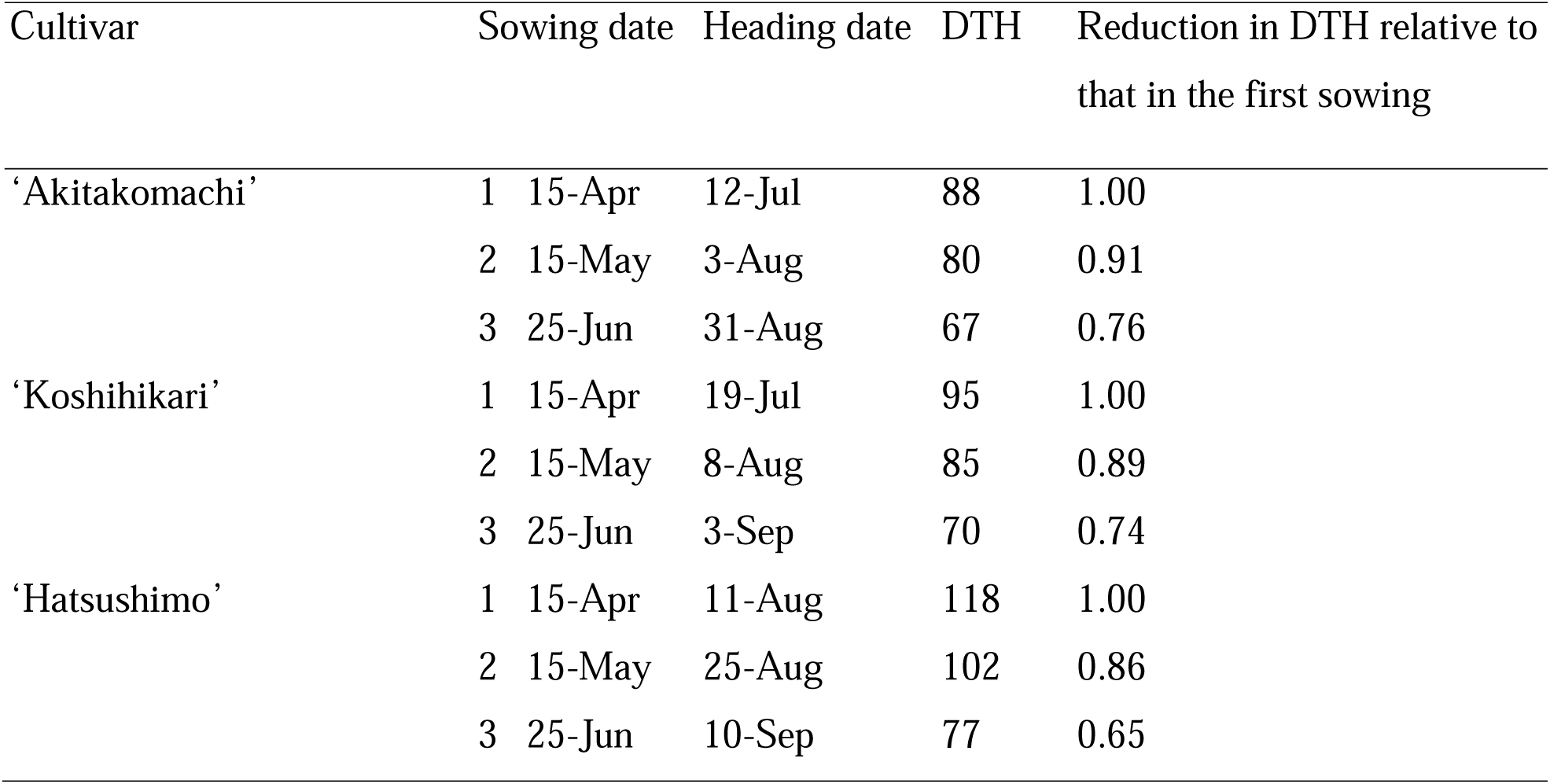
Sowing date, heading date, and days from sowing to heading date (DTH) of the three study cultivars.

Plants were then grown outdoors with the soil surface kept submerged until the start of heat treatments. Tillers emerging during the vegetative stage were removed to ensure uniform panicle development on the main culms. Before heat treatments, 36 pots containing uniform plants were randomly divided into three treatment groups (12 pots per treatment) for each sowing date and cultivar.

### High-temperature treatments

Heat treatments were conducted in three growth chambers (LPH-500-LC, Nippon Medical & Chemical Instruments Co., Ltd., Osaka, Japan), one per treatment. The rice plants were transferred to the chambers at the onset of flowering and exposed for 3 consecutive days to one of three daytime temperature conditions. Diurnal temperature and humidity patterns were set to simulate conditions observed in hot paddy fields in 2020, a hot year in Gifu Prefecture. Daily maximum temperatures were 35, 37, and 39 °C, maintained from 13:00 to 16:00, while nighttime temperatures (18:00–06:00) were set at 24–26 °C in all treatments. Relative humidity was maintained at 60% during the daytime and 90% at night. Temperature and humidity were adjusted stepwise between day and night. Figure 1 shows time profiles of the three treatments.

**Fig. 1.**
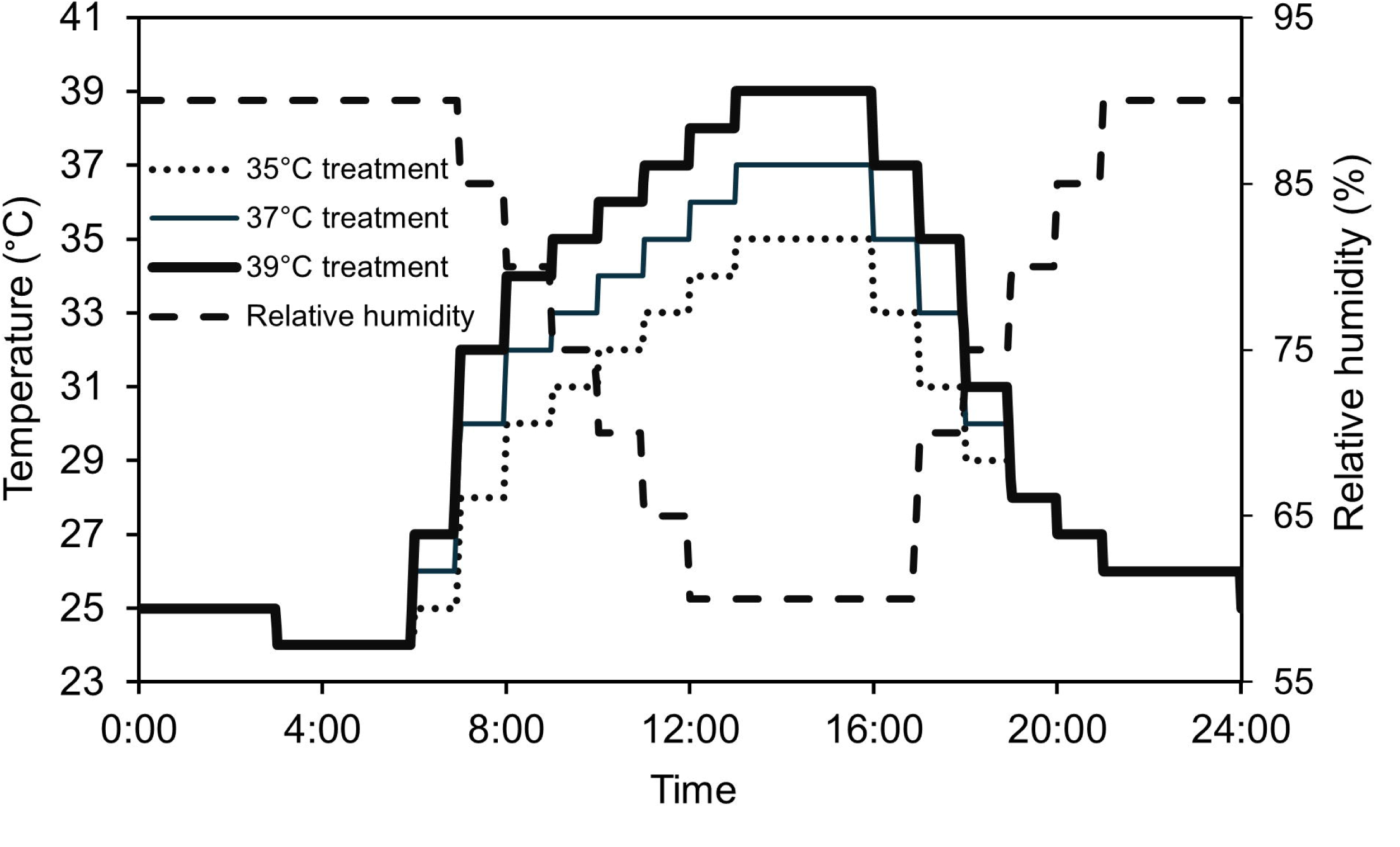
Diurnal changes of air temperatures and relative humidity set in the growth chambers for the three high-temperature treatments.

Temperature and humidity were monitored at panicle height using sensors (HMP45A, Vaisala, Helsinki, Finland) installed in ventilated double cylinders (length 45 cm, inner diameter 7 and 4 cm, airflow 2.15 m s□¹) placed on both the upwind and downwind sides of the plant canopy. Measurements were recorded every minute using data loggers (GL200 and GL240, Graphtec Corporation, Yokohama, Japan). Chamber settings were adjusted to maintain target temperatures on the basis on monitored values.

During the overall treatment period (12 July – 13 September), hourly mean daytime temperatures deviated by +1.17 to −1.83 °C and nighttime temperatures by +1.03 to −1.58 °C from set points. Hourly mean relative humidity deviations ranged from +21.0% to −17.0% during the day and +27.4% to −18.0% at night. During the maximum temperature period (13:00–16:00), mean temperatures deviated by +0.81 to −0.91 °C from target values. Photon flux density and photosynthetic photon flux density at the chamber base (without plants) during daytime (06:00–18:00) were ∼203 and ∼186 µmol m□² s□¹, respectively. Pots were transferred into and out of the chambers at around 18:00.

### Measurements

#### Fertility

Each panicle was tagged with its heading date. Panicles that emerged 1 day before the start of the heat treatment were used for seed set evaluation. These panicles completed most of their flowering during the 3-day treatment period. Seed set was determined by manual inspection of ovarian development at maturity. The rate in each panicle was calculated as the number of florets with swollen ovaries divided by the total number of florets × 100, and the mean value of panicles within a pot was used as the replicate score. At least one panicle per pot was examined. All 12 pots were used for fertility test as replicates.

#### Pollination and anther morphology

During the treatment period, florets opening on a given day were collected after 16:00 from panicles not used for seed set evaluation. Six florets per pot were randomly sampled from eight pots per treatment; four florets were used for pollination observation and the remaining two for anther morphology.

For pollination assessment, stigmas were collected on the day of flowering and stained with cotton blue solution. The numbers of germinated and total (germinated + non-germinated) pollen grains on each stigma were counted under an optical microscope (BX50, Olympus, Tokyo, Japan). The mean value from four florets per pot was used as one replicate. As in previous studies, successful fertilization in rice required either ≥10 germinated pollen grains or ≥20 total pollen grains per stigma (Satake and Yoshida, 1978; Matsui *et al*., 2001). Therefore, the proportions of florets meeting these criteria (GP10 and TP20) were used as indices of pollination stability.

The two florets for anther observation were air-dried in Petri dishes. Anther length and the lengths of apical (ADL) and basal (BDL) dehiscences were measured (Fig. 2) under a digital microscope (KH-7700, Hirox, Tokyo, Japan). Two anthers per floret were measured, and the mean of four anthers was used as one replicate score. Eight scores from eight pots were used as replicates for pollination indices and anther morphologies.

**Fig. 2.**
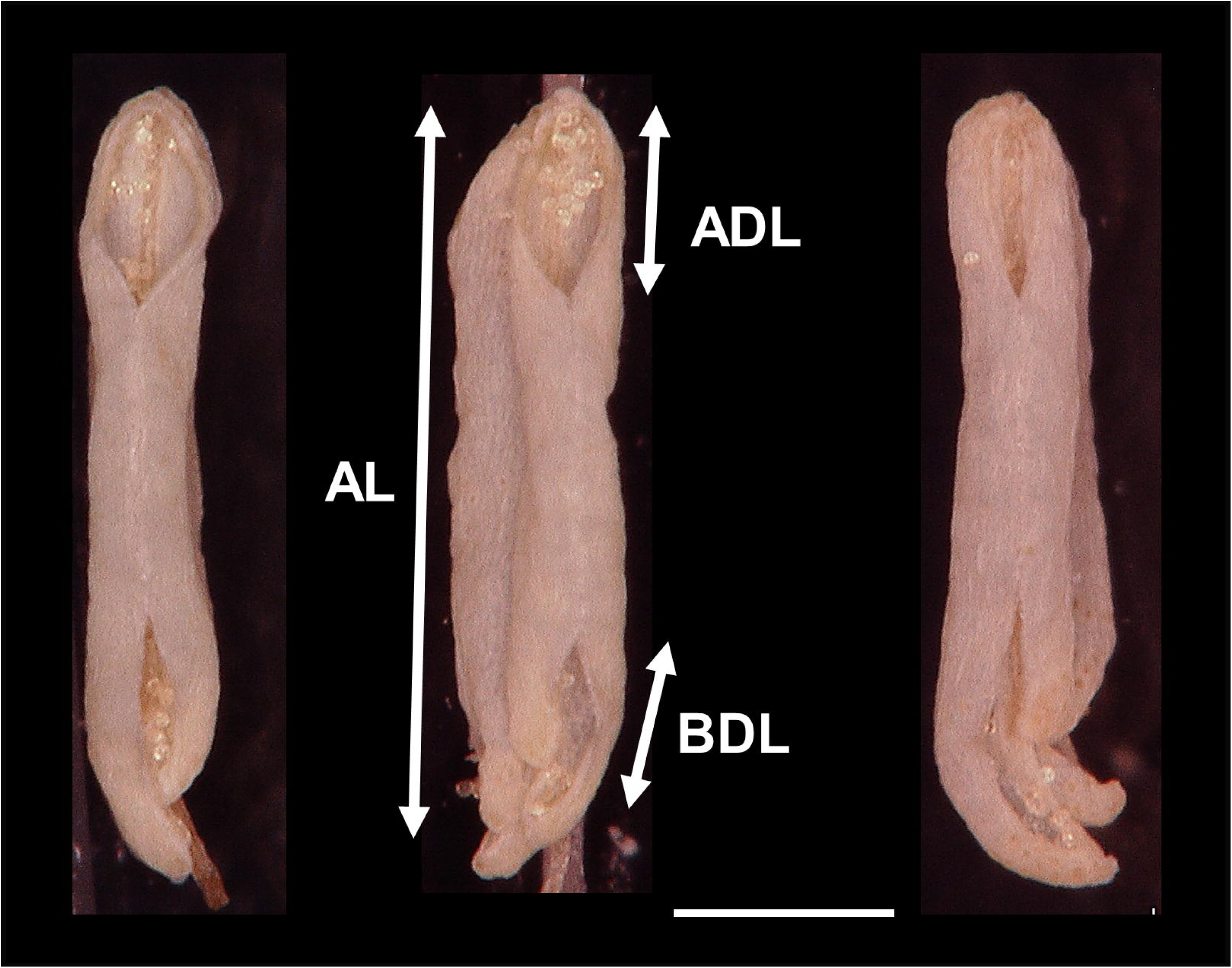
Exterior views of dehisced rice anthers (‘Hatsushimo’) after exposure to high temperature (39 °C) conditions on (left) first, (middle) second, and (right) third sowing dates. Anther length (AL), apical dehiscence length (ADL), and basal dehiscence length (BDL) were measured under a digital microscope. Scale bar, 500 µm.

### Statistical analysis

A four-factor experimental design was used to evaluate the effects of cultivar, sowing date, and temperature treatment on anther morphology, pollination stability, and tolerance to heat-induced sterility. Analysis of variance (ANOVA) was conducted to test the main effects and their interactions, with all factors treated as fixed. When ANOVA results were significant, Tukey’s honestly significant difference (HSD) test was applied for multiple comparisons at *P* < 0.05. Pearson correlation analyses were then conducted for GP10 and floret fertility (Fert) against potentially related variables, including BDL, treatment temperature (for GP10 only), and GP10 and treatment for logit(Fert).

All analyses were conducted in Statistica v. 10 (StatSoft Inc., Tulsa, OK, USA). Fert, GP10, and TP20 were logit-transformed before analysis (logit(FS), logit(GP10), logit(TP20)); an empirical logit transformation was used because FS, GP10, and TP20 include zero values.

## Results and Discussion

### Heading date

Days to heading was shortest in ‘Akitakomachi’ and longest in ‘Hatsushimo’ (Table 1). The relative reduction in days to heading with later sowing date, compared with the first sowing date, was smallest in ‘Akitakomachi’ and largest in ‘Hatsushimo’. This indicates that ‘Hatsushimo’ had the strongest photoperiodic response to sowing date, and ‘Akitakomachi’ had the weakest.

### Anther morphology

ANOVA in anther morphology revealed that the effects of genotype, sowing date, and temperature treatment were all significant for AL and ADL, and the effects of genotype and sowing date were significant for BDL (Table 2). Significant interactions between genotype and sowing date were detected for AL, ADL, and BDL, and between sowing date and treatment for BDL, indicating that the effects of sowing date on anther morphology varied among the cultivars.

**Table 2.**
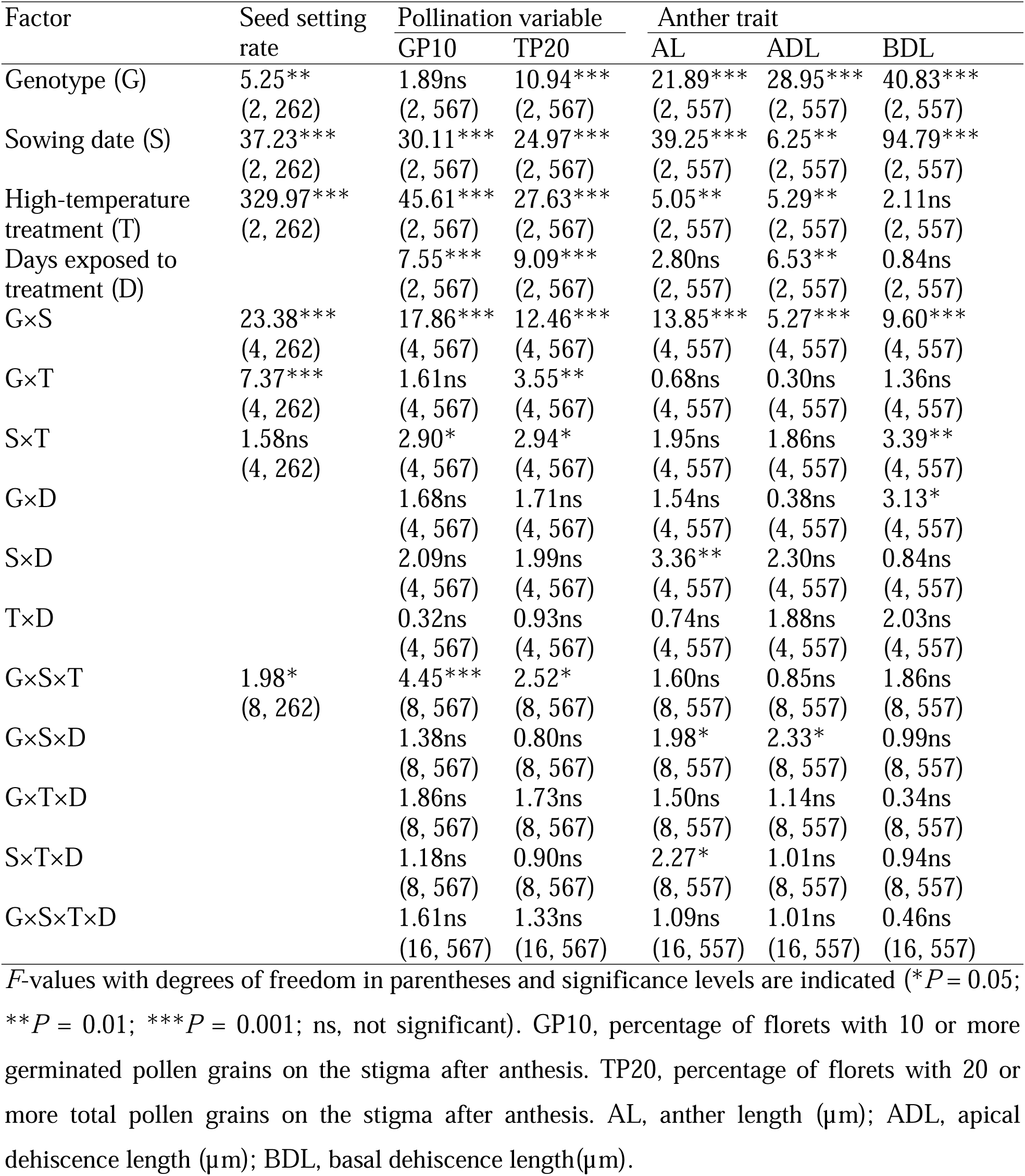
Summary of ANOVA results for the effects of genotype, sowing date, treatment, and duration of exposure to treatment on seed set, pollination-related variables, and dehisced anther traits.

AL tended to decrease with later sowing date, although in ‘Hatsushimo’, AL under the latest sowing recovered to a level comparable to that under the earliest sowing (Table 3). The range of AL among cultivars was relatively narrow (25.7, 51.5, 158.2 μm across the three sowing dates), and the range among sowing dates was similar (across the three cultivars (107.6–131.4 μm). Okamoto and Hara (2021) reported that delayed sowing shortened anthers across nine cultivars on Ishigaki Island, Okinawa, Japan (24°38′ N, 124°20′ E), and that the ratio of days to heading between early and late sowing correlated with AL reduction. This tendency for anthers to shorten with delayed sowing is consistent with these earlier findings. However, we detected no correlation between the relative reduction of days to heading and AL among the three cultivars examined. This discrepancy may reflect differences in pre-flowering growth conditions: our plants were grown outdoors, whereas those of Okamoto and Hara (2021) were grown under controlled temperatures. Environmental conditions before flowering, such as temperature, may therefore influence anther development. Because anther length is correlated with pollen number (Suzuki, 1981) and with tolerance to cool injury (Hashimoto, 1961), environmentally induced modulation of anther length should be an important consideration when evaluating cool-tolerant rice resources.

**Table 3.** Anther length (AL) and anther dehiscence length (µm) at the apical (ADL) and basal (BDL) parts of the anther for the three sowing date and three cultivars.

| Item | Sowing date | Cultivar |  |  | Mean | CV (%) | Range |
| --- | --- | --- | --- | --- | --- | --- | --- |
|  |  | ‘Akitakomachi’ | ‘Koshihikari’ | ‘Hatsushimo’ |  |  |  |
| AL | 1 | 1842.3<sup>c</sup> | 1836.3<sup>b</sup> | 1887.8<sup>b</sup> | 1855.5 | 1.5 | 51.5 |
|  | 2 | 1783.2<sup>b</sup> | 1805.9<sup>b</sup> | 1780.2<sup>a</sup> | 1789.8 | 0.8 | 25.7 |
|  | 3 | 1725.0<sup>a</sup> | 1704.9<sup>a</sup> | 1863.1<sup>b</sup> | 1764.3 | 4.9 | 158.2 |
|  | Mean | 1783.5 | 1782.4 | 1843.7 |  |  |  |
|  | CV (%) | 3.3 | 3.9 | 3.1 |  |  |  |
|  | Range | 117.3 | 131.4 | 107.6 |  |  |  |
| ADL | 1 | 387.8<sup>a</sup> | 439.4<sup>b</sup> | 429.0<sup>b</sup> | 418.7 | 6.5 | 51.6 |
|  | 2 | 393.2<sup>a</sup> | 392.2<sup>a</sup> | 421.9<sup>b</sup> | 402.5 | 4.2 | 29.7 |
|  | 3 | 392.4<sup>a</sup> | 426.0<sup>b</sup> | 429.8<sup>b</sup> | 416.0 | 5.0 | 37.4 |
|  | Mean | 391.1 | 419.2 | 426.9 |  |  |  |
|  | CV (%) | 0.7 | 5.8 | 1.0 |  |  |  |
|  | Range | 5.4 | 47.2 | 7.9 |  |  |  |
| BDL | 1 | 423.9<sup>b</sup> | 456.7<sup>a</sup> | 479.0<sup>b</sup> | 453.2 | 6.1 | 55.1 |
|  | 2 | 383.1<sup>a</sup> | 445.8<sup>a</sup> | 403.4<sup>a</sup> | 410.8 | 7.8 | 62.7 |
|  | 3 | 451.1<sup>b</sup> | 551.6<sup>b</sup> | 588.2<sup>c</sup> | 530.3 | 13.4 | 137.1 |
|  | Mean | 419.4 | 484.7 | 490.2 |  |  |  |
|  | CV (%) | 8.2 | 12.0 | 19.0 |  |  |  |
|  | Range | 68.0 | 105.8 | 184.8 |  |  |  |

Within columns, values sharing the same letters are not significantly different by Tukey’s HSD test.

For ADL, no effect of sowing date was observed in ‘Akitakomachi’ and ‘Hatsushimo’, but ADL was significantly shorter under the second sowing date than under the other sowing dates in ‘Koshihikari’ (Table 3). The ranges and coefficients of variation for AL and ADL were all smaller than those for BDL, indicating greater plasticity in basal dehiscence than in other traits. The range of mean BDL across sowing dates (68.0–184.8 µm in three cultivars) was comparable to or greater than that across cultivars (55.1–137.1 µm in three sowing dates), suggesting that environmental conditions associated with sowing date exerted a stronger influence than genetic differences on BDL. Across the sowing dates, all cultivars exhibited the shortest BDL under the second sowing date and the longest under the third sowing date, with a significant difference between them (Table 3). ‘Hatsushimo’ showed the greatest BDL at the third sowing date and the largest variation across sowing dates among cultivars.

Overall, the influence of sowing date on BDL was most pronounced in the late-heading ‘Hatsushimo’. Differences in heading date are associated with variation in environmental conditions during growth and with genetic differences in photoperiod sensitivity, both of which may affect BDL development.

### Pollination stability and fertility

For GP10, the effects of temperature treatment and sowing date were significant, whereas the effect of genotype was not (Table 2). Significant interactions were detected for genotype × sowing date, sowing date × treatment, and genotype × sowing date × treatment, indicating complex genotype–environment interactions in pollination stability.

In ‘Akitakomachi’, GP10 was significantly higher under the first sowing date than under the second and/or third sowing dates (Table 4). In ‘Koshihikari’, GP10 was also significantly higher under the first sowing date than that under the second and third sowing dates in all three treatment temperatures. In ‘Hatsushimo’, GP10 was significantly higher under the third sowing date than under the first sowing date in the 35 °C treatment, but there were no significant differences in the other two treatments. Because the interaction between treatment and sowing date was not significant in ‘Hatsushimo’ (data not shown), multiple comparisons among sowing dates were conducted using mean values for the three temperature treatments, and significant differences were detected between GP10 under the third sowing date and that under the other sowing dates (Table 4).

**Table 4.**
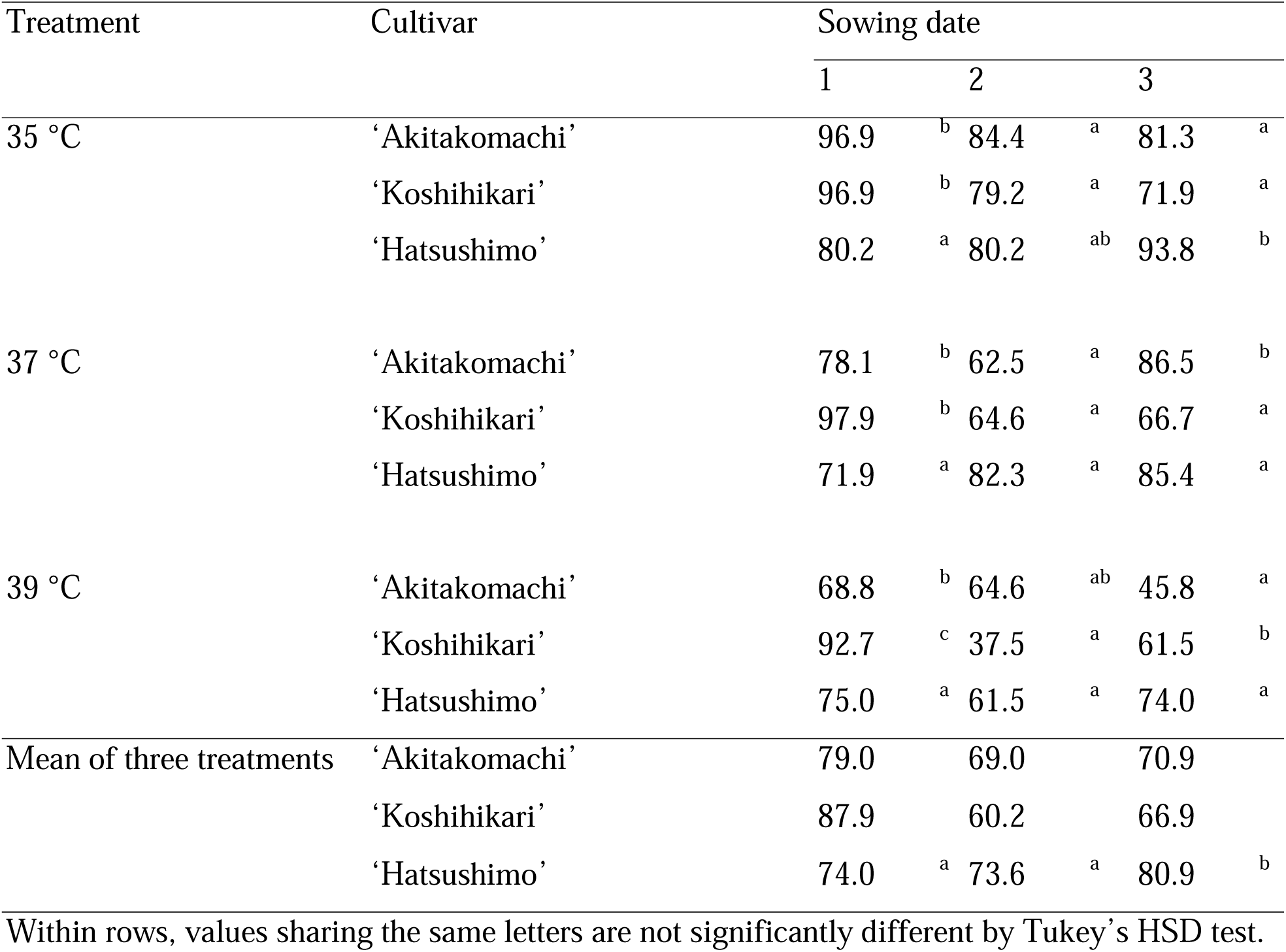
Effect of sowing date on logit-transformed GP10 (% of florets with ≥10 germinated pollen grains on the stigma after anthesis) across all treatment and cultivar combinations.

For fertility rate, the effects of genotype, sowing date, and temperature treatment were all significant. Significant interactions were also observed for genotype × sowing date, for genotype × treatment, and for genotype × sowing date × treatment (Table 2), again indicating complex genotype–environment effects on fertility.

In ‘Akitakomachi’ and ‘Koshihikari’, fertility rate was significantly lower under the second sowing date than under the other dates, except in ‘Koshihikari’ at 35 °C and ‘Akitakomachi’ at 39 °C, where differences across sowing dates were not significant. In these exceptions, however, the mean values under the second sowing date were still the lowest among the three sowing dates (Table 5). In contrast, in ‘Hatsushimo’, fertility was significantly lower under the first sowing date than under the third sowing date in all temperature treatments. These results suggest that the second sowing date was unfavorable for fertility under heat stress in ‘Akitakomachi’ and ‘Koshihikari’, and the first sowing date was unfavorable in ‘Hatsushimo’.

**Table 5.**
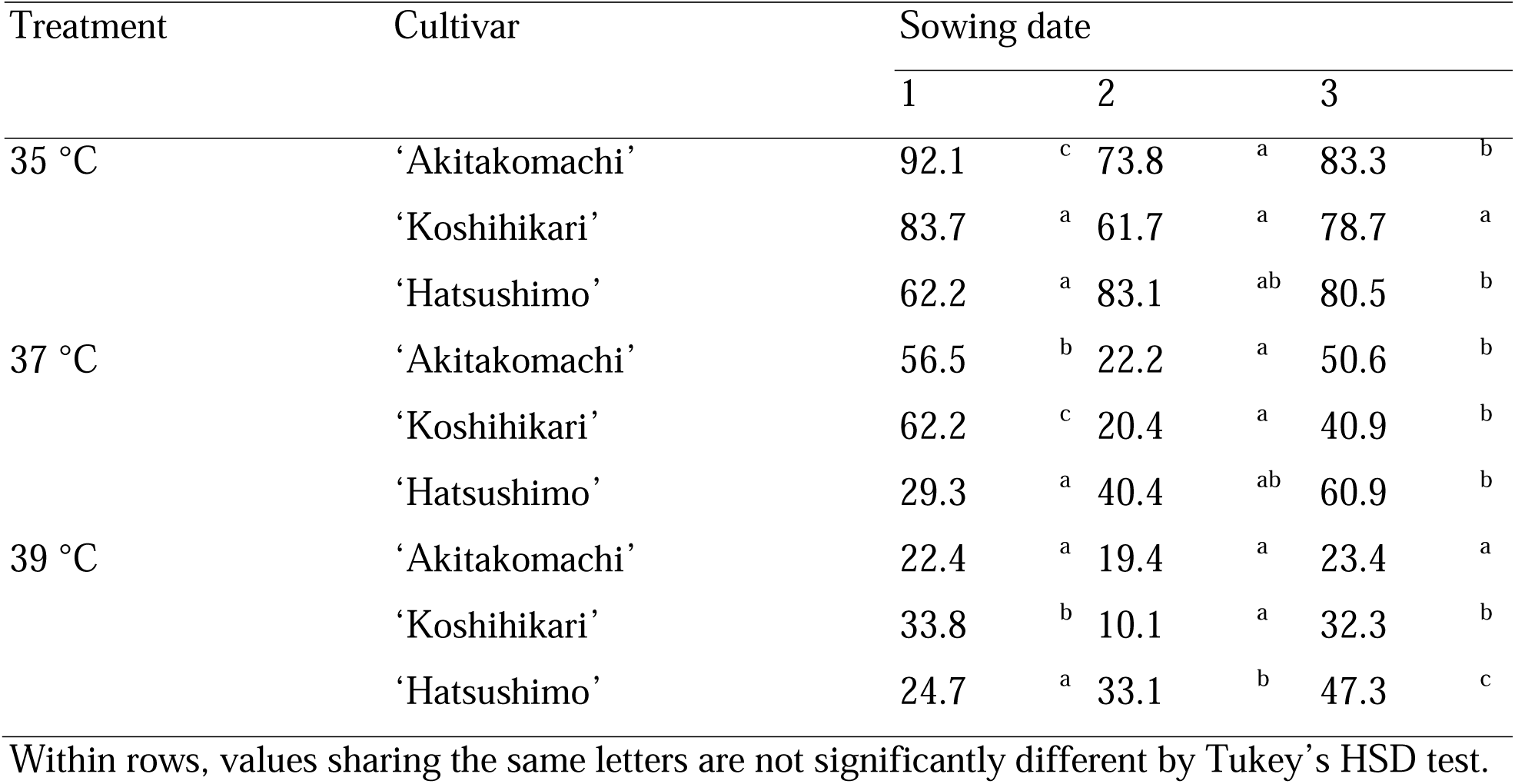
Effect of sowing date on seed set across all treatment and cultivar combinations.

### Factors determining fertility among cultivars

A long BDL promotes stable pollination (Matsui and Kagata, 2003; Tazib *et al*., 2015). Here, both genotype and sowing date significantly affected BDL, and their interaction was complex. Therefore, to clarify the effects of sowing date mediated through BDL, we analyzed each cultivar individually. Because daytime chamber temperatures could not be strictly controlled during the unusually hot summer of 2024, we used actual measured temperatures in the analyses. For example, in the 39 °C treatment under the third sowing date in ‘Hatsushimo’, the actual mean temperature between 13:00 and 16:00 was 1.4 °C higher than that at the first sowing date. As a 1-°C difference in temperature at flowering can alter fertility by 10%–20% (Matsui *et al*., 2001), we used the measured maximum temperature during 13:00–16:00 instead of the chamber setpoint in the initial regression analyses.

For ‘Akitakomachi’ and ‘Koshihikari’, logit(GP10) was well explained by maximum temperature alone (Equation 1 and 2, Table 6):

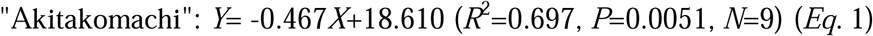

**Table 6.** Summary of regression analysis results for logit(GP10).

| Cultivar | Explanatory variable | Regression coefficient, B*, or constant | Standardized regression coefficient or $\beta^{**}$ | Partial correlation coefficient | *P* | *R*<sup>2</sup> |
| --- | --- | --- | --- | --- | --- | --- |
| ‘Akitakomachi’ | Temp | −0.467 | −0.835 | – | 0.005 | 0.697 (*N* = |
|  | BDL | – | – | – | ns | 9, *P* = |
|  | Intercept | 18.610 |  |  | 0.004 | 0.0051) |
| ‘Koshihikari’ | Temp | −0.310 | −0.668 | – | 0.049 | 0.446 (*N* = |
|  | BDL | – | – | – | ns | 9, *P* = |
|  | Intercept | 12.417 |  |  | 0.038 | 0.0493) |
| ‘Hatsushimo’ | Temp | −0.193 | −0.756 | −0.694 | 0.012 | 0.739 (*N* = |
|  | BDL | 2.671 | 0.576 | 0.652 | 0.035 | 9, *P* = |
|  | Intercept | 7.030 |  |  | 0.012 | 0.0178) |

\*Partial regression coefficient
\*\*Standardized partial regression coefficient

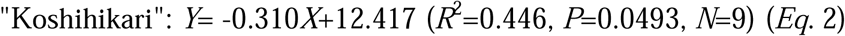

where *Y* is logit(GP10) and *X* is maximum temperature (°C). In contrast, in ‘Hatsushimo’, both maximum temperature (*X*□) and BDL (*X*□) significantly explained logit(GP10) (Equation 3):

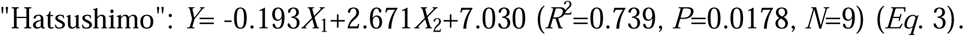

The statistical significance for BDL in the absence of genetic variation suggests that environmental effects associated with sowing date influenced logit(GP10) through variation in BDL.

It has been reported that more than 10 germinated pollen grains on the stigma are required for successful fertilization in rice (Satake and Yoshida, 1978). Indeed, GP10 and fertility have shown generally good agreement under both controlled environmental (Matsui *et al*., 2001) and field conditions (Matsui *et al*., 2021, 2025), except at high temperatures (>35 °C). Here, we also found a significant correlation between logit(GP10) and logit(Fert) across all genotypes and treatments:

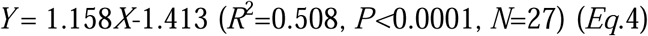

where Y is logit(Fert) and X is logit(GP10). The product of the regression slopes of the relationships between BDL and logit(GP10) (2.671) and between logit(GP10) and logit(Fert) (1.158) was 3.093, close to the partial regression coefficient (2.7) reported under field conditions in China (Matsui *et al*., 2021). This suggests that a 0.1-mm increase in BDL could improve GP10 by ∼15% (at GP10 ≈ 50%) and seed set (FS) by ∼6.7% (at FS ≈ 50%), consistent with previous field studies (Tian *et al*., 2010; Zhao *et al*., 2010).

Under extremely hot conditions (∼40 °C), flowering time and daily maximum temperature can directly reduce fertility, not only via pollen germination on the stigma (Matsui *et al*., 2001, 2021). Here, the effect of heat treatment on logit(Fert) was also significant (Fig. 3). As treatment temperature increased, logit(Fert) declined relative to logit(GP10). When temperature treatment was included in the model, logit(GP10) closely approximated logit(Fert) (*R*² = 0.784, *P* < 0.0001; Fig. 3). This indicates that at temperatures between 35 and 39 °C, higher levels of pollen germination are required for successful fertilization, and increases in logit(GP10) remain effective in improving logit(Fert), consistent with field observations (Matsui *et al*., 2021, 2025).

**Fig. 3.**
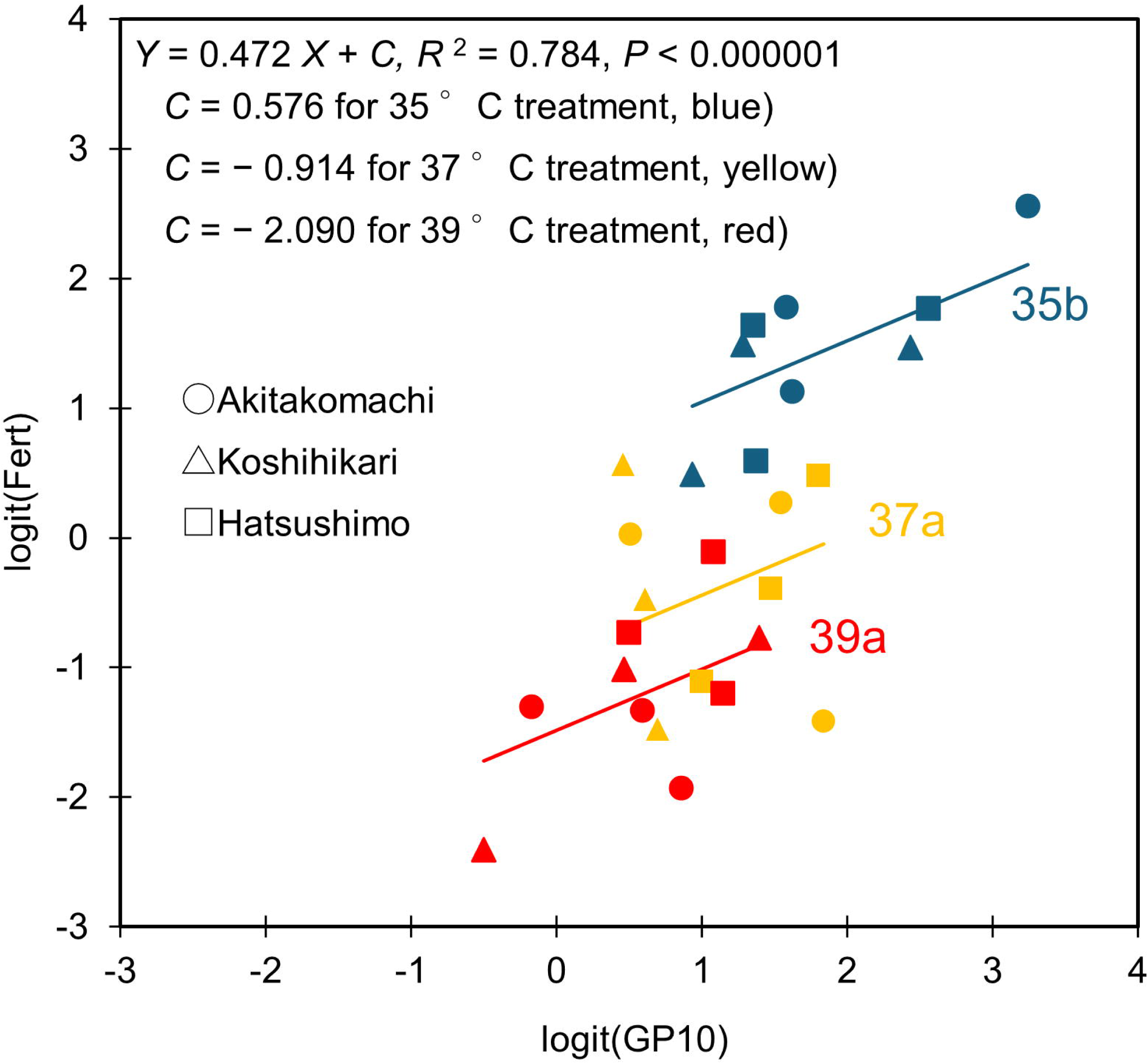
Relationship between logit(GP10) and logit(Fert) for each of the study cultivars under the three high-temperature treatment conditions. Analysis of covariance, with treatment as a fixed variable and logit(GP10) as a covariate, showed significant effects of both treatment (*P* < 0.01) and logit(GP10) (*P* < 0.05) on Logit(Fert). Lines sharing the same letters are not significantly different. *R*^2^, coefficient of determination for the model.

The standardized partial regression coefficient of BDL for logit(GP10) was 0.576 in the multiple regression analysis (Equation 3, Table6) and that of logit (GP10) for logit (Fert) was 0.290 in the covariance analysis (data not shown). Thus, the cumulative effect of BDL on logit(Fert), viewed through path analysis, was 0.167, close to the correlation coefficient (0.175) between BDL and logit(Fert) in ‘Hatsushimo’ (data not shown). Because logit(GP10) was correlated with temperature at flowering (data not shown), multicollinearity may have affected the relationships among logit(GP10), temperature, and logit(Fert). Nevertheless, the explanatory model provided a good approximation.

Overall, these results demonstrate strong genotype × environment interactions in determining anther morphology and fertility. In rice, anther dehiscence is a key determinant of tolerance to high-temperature–induced sterility through its effect on pollination stability (Matsui *et al*., 2021). Because anther dehiscence can be easily observed microscopically and correlates between normal and hot conditions (Matsui *et al*., 2005), it is a potentially useful visual marker for selecting heat-tolerant lines. However, our results highlight three important caveats: (i) anther dehiscence size varies with both growth environment and genotype, (ii) environmental effects on anther dehiscence differ among genotypes, and (iii) environmentally induced variation can be comparable in magnitude to genetic differences. Therefore, when cultivars are selected on the basis of anther dehiscence traits, environmental differences among locations and cropping seasons must be carefully considered to ensure reliable assessment.

### Conclusion

Both genotype and pre-flowering environmental conditions substantially influenced anther morphology, pollination stability, and fertility in rice under high temperatures. Among the anther traits examined, BDL showed the greatest environmental variability. BDL affected logit(GP10) in the late-heading ‘Hatsushimo’, and regression analyses indicated that the BDL affects fertility primarily through its effect on the number of germinated pollen grains on the stigma.

Although anther dehiscence, especially BDL, can serve as a practical morphological marker for selecting heat-tolerant genotypes, our findings highlight two important cautions: (i) BDL is environmentally plastic during plant growth, and (ii) the magnitude of the environmental response differs among genotypes. Therefore, evaluation and selection of heat-tolerant rice cultivars on the basis of anther traits should carefully consider environmental differences among locations and cropping seasons to ensure reliable assessment.

## Acknowledgements

We thank the staff of the Faculty of Applied Biological Sciences, Gifu University, Japan, for maintaining the experimental facilities.

## Author contributions

KK: investigation, formal analysis, data curation, writing—original draft, visualization; TY: writing—review & editing, supervision; TM: conceptualization, methodology, formal analysis, data curation, resources, writing—review & editing, visualization, supervision.

## Conflict of interest

No conflict of interest declared.

## Funding

This research received no specific grant from any funding agency in the public, commercial or not-for-profit sectors.

## Data availability

The datasets generated and/or analyzed during the current study are available from the corresponding author on reasonable request.

## Abbreviations

AL (mm or μm): anther length
ADL (mm or μm): length of dehiscence for pollen dispersal formed at the apical part of the anther
BDL (mm or μm): length of dehiscence for pollen dispersal formed at the basal part of the anther
FS (%): percentage of floret sterility
GP10 (%): percentage of florets with 10 or more germinated pollen grains on the stigma after anthesis
TP20 (%): percentage of florets with 20 or more total pollen grains on the stigma after anthesis

## Notes

### Competing Interest Statement

The authors have declared no competing interest.

